# PIEZO channels link mechanical forces to uterine contractions in parturition

**DOI:** 10.1101/2025.09.17.676899

**Authors:** Yunxiao Zhang, Sejal A. Kini, Sassan A. Mishkanian, Renhao Luo, Saba Heydari Seradj, Verina H. Leung, Yu Wang, M. Rocío Servín-Vences, William T. Keenan, Utku Sonmez, Oleg Yarishkin, Manuel Sanchez-Alavez, Yuejia Liu, Xin Jin, Darren J. Lipomi, Li Ye, Michael Petrascheck, Antonina I. Frolova, Sarah K. England, Ardem Patapoutian

## Abstract

Mechanical forces are extensively involved in pregnancy and parturition, but their precise roles and mechanisms remain poorly understood. Here, we identify mechanically activated ion channels PIEZO1 and PIEZO2 as key mechanotransducers required for labor progression. Genetic deletion of *Piezo1* and *Piezo2* in mice resulted in weakened uterine contractions and severe parturition defects. Tissue-specific knockouts revealed that deletion in either the uterus or sensory neurons alone caused modest defects, whereas combined loss significantly impaired labor, demonstrating additive effects. Single-nuclei sequencing showed that loss of PIEZO reduced expression of connexin43 (*Gja1*), a gap junction protein in uterine smooth muscle cells, suggesting a mechanistic link to impaired contraction. These findings highlight the critical role of PIEZO channels in mechanotransduction during parturition and suggest therapeutic targets for labor dysfunction.

---

Mechanotransduction, the cellular response to mechanical stimuli, is essential for a wide range of biological processes, from tactile perception to the maintenance of tissue homeostasis (*1–4*). Although both mechanical cues and chemical ligands influence physiological functions across multiple organ systems, including cardiovascular (*5*), respiratory (*6*) and gastrointestinal activities (*7*), the extent to which mechanical forces contribute under different physiological conditions remains less well characterized.

Pregnancy and parturition are key physiological events in which mechanical forces play a critical role alongside hormonal and other biochemical signals. During pregnancy, the human uterus can increase its volume capacity by up to 500-fold (*8*), and parturition requires coordinated uterine contractions and extensive cervical remodeling—processes inherently accompanied by dramatic changes in mechanical forces (*9*). Although hormones, particularly progesterone, are well-documented regulators of pregnancy (*10, 11*), evidence indicates that mechanical cues also play a critical and complementary role. As early as 1941, studies by Ferguson et al., suggested that mechanical distention at the vagina stimulates uterine contractions as parturition approaches (*12*). Since then, accumulating evidence has linked mechanical cues to uterine hypertrophy, myometrial contractility, and cervical softening (*9, 13*), all of which are essential for successful reproduction in mammals.

Multiple signaling mechanisms —including mitogen-activated protein kinase (MAPK) signaling (*14, 15*), Hippo/YAP signaling (*16*), matrix metalloproteinases (MMP) (*17*), and ion channels (*18–20*)—have been implicated in mechanosensation. However, since most of these pathways are also activated by biochemical cues, it remains challenging to determine whether the effects of disrupting these pathways are solely due to the loss of mechanotransduction. In contrast, PIEZO1 and PIEZO2 are directly activated by mechanical forces, and unlike many other signaling pathways, lack known endogenous chemical ligands that modulate their activity (*21*). Thus, phenotypes associated with PIEZO knockdown implicates a role of mechanosensation. Here we demonstrate that PIEZO1 and PIEZO2 are specifically required for effective uterine contractions and successful parturition. Knockout of these channels result in significantly impaired uterine contractions and prolonged labor, suggesting mechanotransduction as a critical regulatory mechanism for parturition.

## Results

### Distinct expression patterns of *Piezo1* and *Piezo2* in the reproductive system

We first characterized the distribution of *Piezo1* and *Piezo2* in the mouse reproductive system using single molecule fluorescence *in situ* hybridization (smFISH). At gestational day 18.5 (GD18.5, ∼1 day before parturition), *Piezo1* mRNA was widely detected across most cell types within the uterine horn, whereas *Piezo2* expression was confined to a small subset of cells within the decidualized stroma (Fig. 1, A and B). A similar expression pattern was observed in non-pregnant uterine horns at estrus (Fig. S1, A and B), a finding corroborated by published single cell sequencing data (*22*). Moreover, the overall expression patterns of both *Piezo1* and *Piezo2* remain largely unchanged throughout different stages of the estrous cycle (Fig. S1 C and D). Notably, among the cell types analyzed, only the lymphatic endothelial cells (LECs), which are primarily localized between circular and longitudinal myometrium at late pregnancy (GD18.5), showed high levels of *Piezo2* expression (Fig. S1E). In the cervix, while *Piezo1* was broadly expressed across various cell types, *Piezo2* expression was predominantly observed in epithelial cells (Fig. S1, F to H).

**Figure 1.**
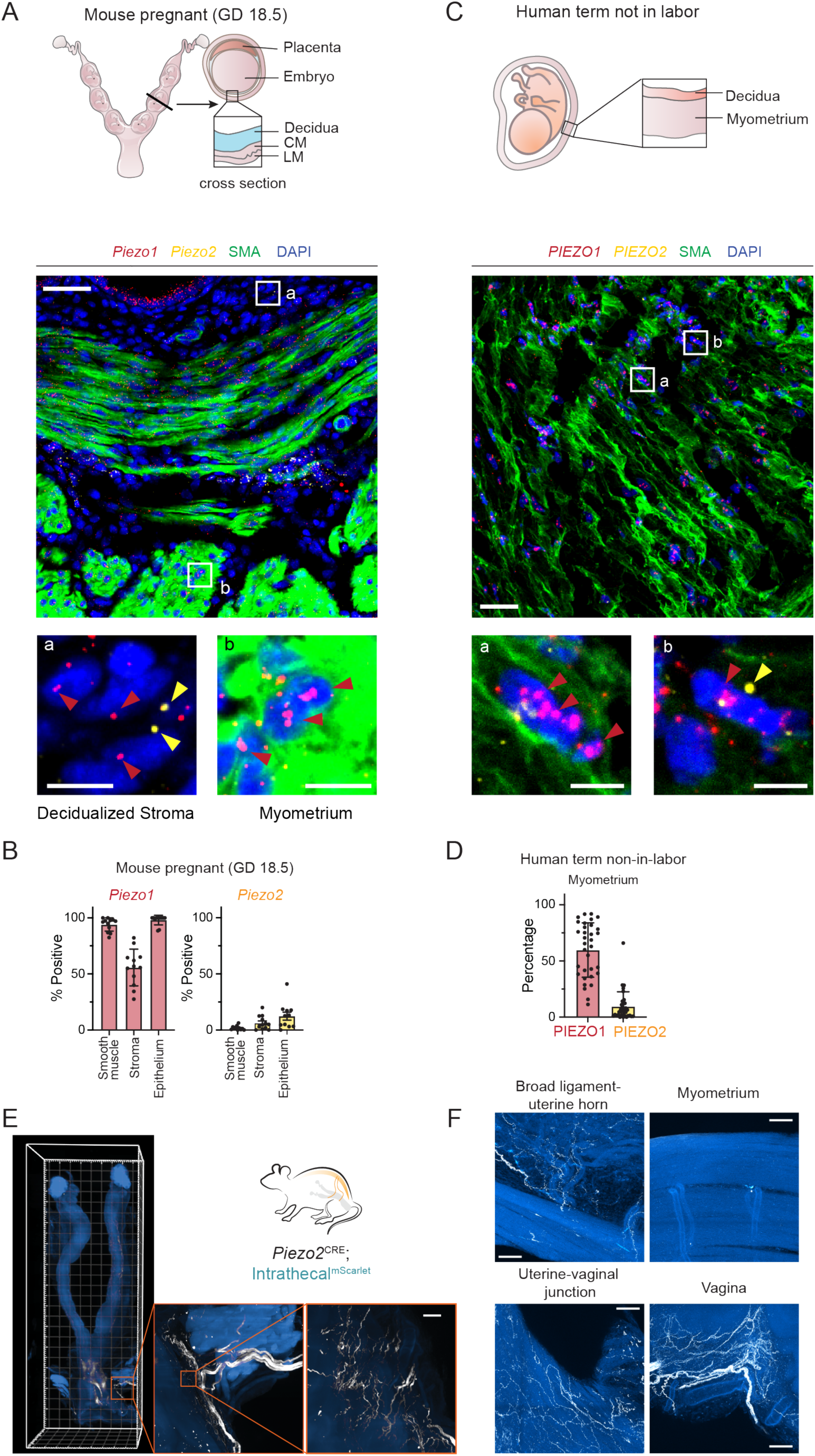
PIEZO1 and PIEZO2 expression in the uterus. (A) At late pregnancy (gestational day 18.5), PIEZO1 is expressed in smooth muscle cells and stromal cells in the uterus, while PIEZO2 is expressed only in a small subset of cells. Scale bar: 50 µm in the overview, 10 µm in the zoomed inset. The percentage of positive cells is quantified in (B). Representative images from 3 animals. CM: circular myometrium. LM: longtitudinal myometrium. (C) Term-not-in-labor human myometrium reveals that PIEZO1 is broadly expressed while PIEZO2 is only sparsely expressed in limited number of cells. Scale bar: 50 µm in the overview, 10 µm in the zoomed inset. The percentage of positive cells is quantified in (D). Representative images from 3 donor specimens. (E) Piezo2-positive sensory innervation is labeled by intrathecal delivery of MacPNS1 AAV virus encoding CAG-DIO-mScarlet into Piezo2-cre animals. The Piezo2 sensory fibers are most abundant in the lower reproductive tract. (F) Confocal imaging revealed Piezo2-positive fibers in the vaginal walls and at the junction between uterine body and vagina but are rare within the uterine horns. Piezo2 fibers are also present in the broad ligament around the uterine horns, and some may go along major vasculature towards the horns. Images were collected from 3 animals. Scale bar: 100 µm

We next assessed *PIEZO1* and *PIEZO2* expression in human reproductive tissues. In non-labor term myometrium, *PIEZO1* was broadly expressed in smooth muscle cells, whereas *PIEZO2* expression was scarce (Fig. 1, C and D). Single cell sequencing data from laboring term myometrium confirmed these observations (*23*) and further indicated that *PIEZO2* expression was restricted to lymphatic endothelial cells (Fig. S1, J and K). Taken together, these findings reveal a remarkable similarity between human and mouse tissues, with widespread PIEZO1 expression and limited PIEZO2 expression.

The reproductive tract receives sensory innervation that may detect mechanical forces such as stretch and pressure within the uterus and cervix (*24*). Since PIEZO2 is abundantly expressed in dorsal root ganglia (DRG) and plays a prominent role in somatosensation and interoception (*25*), we investigated whether it contributes to mechanosensation in the reproductive tract. We labeled *Piezo2^+^* sensory neurons by intrathecally delivering adeno-associated virus (AAV) expressing a Cre-dependent fluorescent protein into *Piezo2*-ires-Cre animals (*Piezo2^Cre^*; intrathecal^mScarlet^). This method efficiently labeled DRGs below the T13 vertebral level. Light sheet microscopy revealed that *Piezo2*^+^ sensory fibers primarily innervate the lower reproductive tract (Fig. 1E). High-resolution confocal imaging further showed abundant *Piezo2*^+^ fibers in the vagina and lower uterus, especially near the cervix (Fig. 1F). In contrast, only sparse *Piezo2*^+^ fibers extended into the uterine horn along the broad ligament vasculature, and the myometrium exhibited almost no labeling (Fig. 1F). Given that nerve fiber density in the uterine horn dramatically decreases during late gestation (*26, 27*), it is expected that *Piezo2*-dependent sensory afferents would originate almost exclusively from the lower reproductive tract at late pregnancy. Moreover, retrograde labeling with cholera toxin subunit B (CTB) revealed minimal overlap with *Sst*, a marker for *Piezo1*^+^ neurons (*28*). Among 240 CTB-labeled neurons from 4 animals (Fig. S1L), no overlap was observed, indicating that *Piezo1*^+^ innervation is rare in reproductive tissues. Taken together, these findings indicate that *Piezo1* is predominantly expressed within reproductive tissues, whereas *Piezo2* primarily marks the sensory afferents.

### *Piezo1* and *Piezo2* knockout mice impair parturition but not gestation

To investigate the functional roles of PIEZO channels during pregnancy and parturition, we generated mice with *Piezo1/2* deletion using *HoxB8^Cre^*, broadly targeting tissues caudal to the diaphragm. *HoxB8^Cre^* covered most cells in the uterus except for the epithelium as indicated by the Sun1 reporter in non-pregnant animals (Fig. S2A).

We further examined the knockout efficiency in uterine horns at GD18.5 by smFISH. *Piezo1* expression was significantly reduced in *SMA*+ (smooth muscle actin) smooth muscle cells within the myometrium of the knockout animals (*HoxB8^Cre^*; *Piezo2 ^f/f^*; *Piezo2^f/f^*, referred to as HoxB8; P1; P2) whereas its expression in adjacent non-smooth muscle cells, which are primarily lymphatic endothelial cells, remained largely unchanged (Fig. S2, B and C).

To avoid potential complications from knocking out PIEZOs in the developing fetuses, we set up the breeding using wildtype C57BL6/J males so that all embryos in gestation maintained at least one copy of functional *Piezo1/2* (Fig. 2A). As previously reported, *HoxB8^Cre^* mediated *Piezo2* knockout exhibited impaired mating behavior, failing to complete intercourse during the standard single-night mating protocol (*29*). Nonetheless, continuous mating attempts eventually resulted in successful pregnancies. The overall appearance and size of the embryos in gestation were comparable to Cre-control littermates (Fig. 2, B and C), indicating minimal involvement of PIEZO1/2 in gestation.

**Figure 2.**
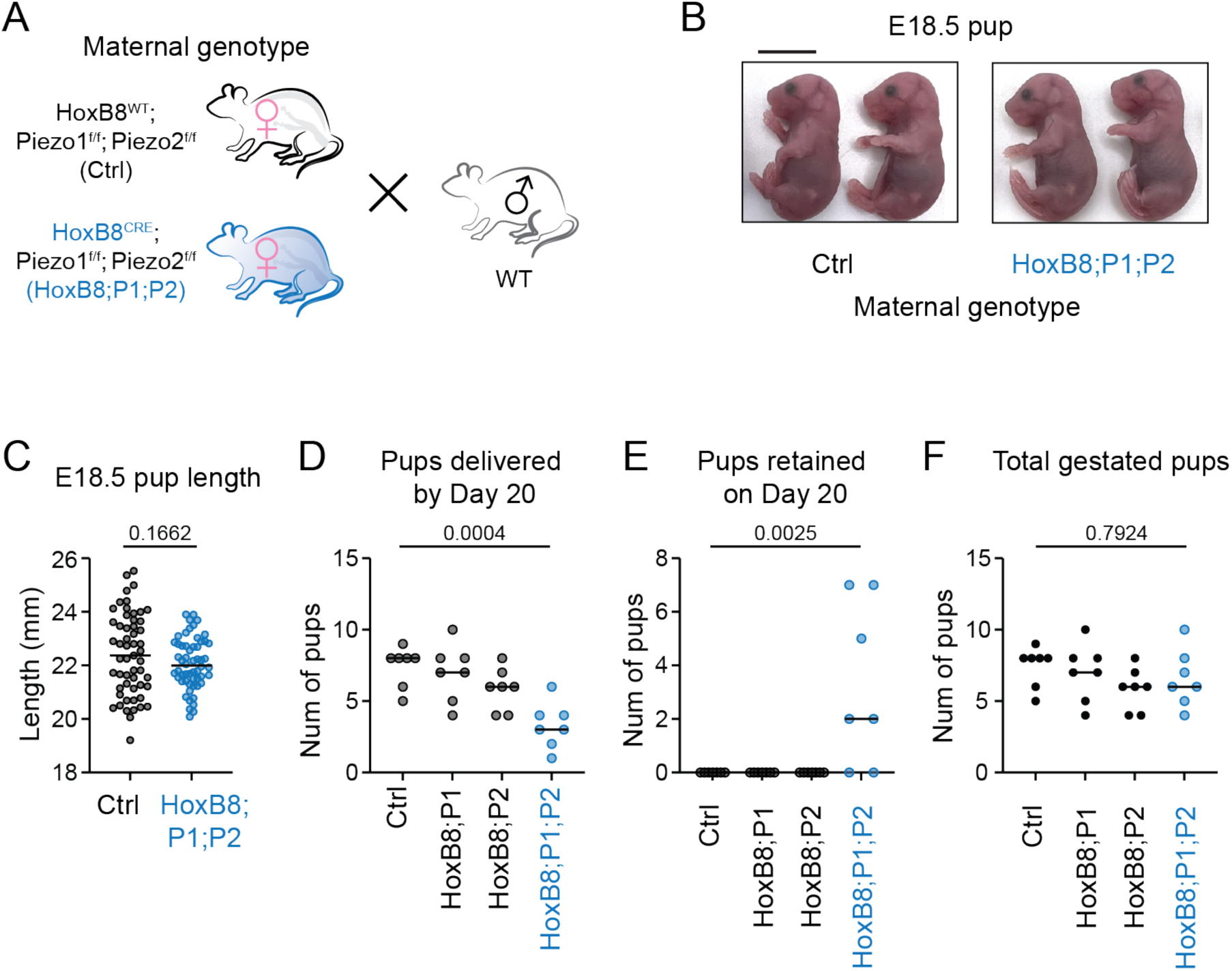
PIEZO1/2 knockout animals have apparently normal gestation but parturition defects. (A) HoxB8 Cre is used to knock out PIEZO1 and PIEZO2 in most cells relevant to pregnancy. The female animals are mated to wild type males to ensure that the offspring have at least one functional copy of PIEZO1/2 and avoid developmental defects. (B) The embryos at gestational day 18.5 share similar appearance between knockout and control animals and the overall dimensions as measured by crown-rump length are within the same range in (C). Statistical test: t-test. Scale bar: 1cm (D) Fewer pups are delivered by PIEZO1/2 double knockout animals by GD 20, as some animals still had pups retained on GD 20 as in (E). The total number of pups in gestation is similar regardless of the genotype as in (F). Deletion of PIEZO1 or PIEZO2 alone does not affect the number of pups delivered by day 20. Brown-Forsythe One-way ANOVA was used for statistical testing.

At parturition, however, significant complications were observed in *Piezo1/2* double-knockout females, characterized by prolonged labor and incomplete delivery by gestational day 20 (GD20) (Fig. 2, D to F). All animals started parturition on GD19, but retained embryos were still present in many double knockout animals by GD20. The total number of embryos in gestation was similar across genotypes, suggesting that *Piezo1/2* did not affect general fertility but had a specific effect on parturition. Single knockouts showed no evident parturition deficits, suggesting compensatory interactions between *Piezo1* and *Piezo2*.

As complications in parturition are frequently attributed to issues in cervix remodeling and uterine contraction, we performed histological analysis on uterus and cervix sections. Uterine horns at GD18.5 from control and knockout animals revealed typical layered structures in hematoxylin-eosin (H&E) staining (Fig. S2D). Cervix sections revealed no discernible abnormalities in extracellular matrix (ECM) remodeling by trichrome staining (Fig. S2E). The parturition defects are thus less likely due to complications with cervical remodeling.

### Loss of *Piezo1* and *Piezo2* impairs uterine contraction

To directly assess uterine contractions, we used implantable pressure sensors to continuously measure intrauterine pressure in free-moving animals during labor. We also used a 3D-printed cage lid equipped with infrared LED lights and camera to monitor animals in their home cages and assess parturition events (Fig. 3, A and B). In this way, we enabled the animals to enter spontaneous labor and obtained data under physiological conditions.

**Figure 3.**
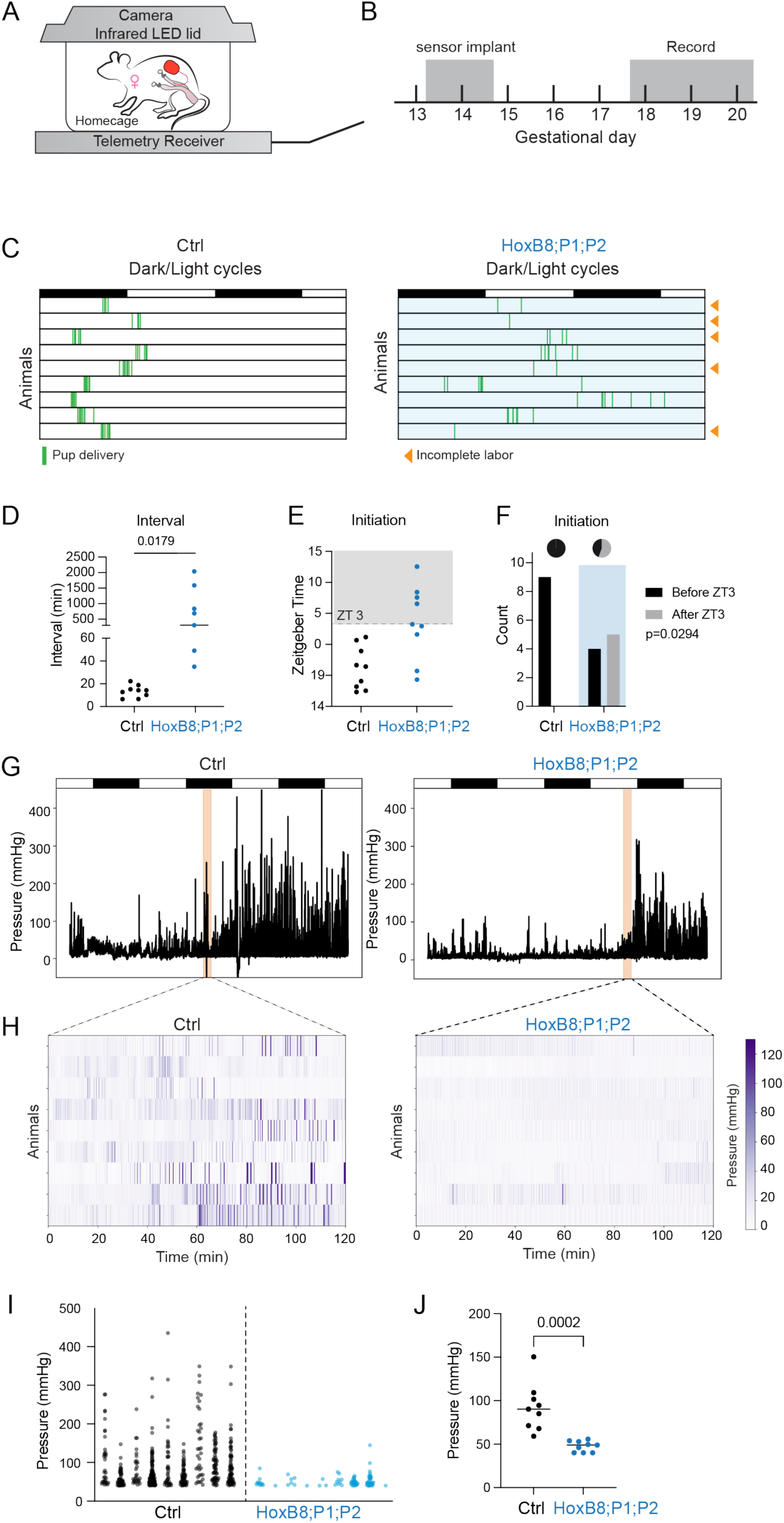
PIEZO1/2 knockout animals have defects in uterine contraction. (A) The recording setup consists of a receiver that records intrauterine pressure signals from the telemetry implants in real time and a custom-made cage lid with infrared LED lights and a camera to record animal activity. (B) Telemetry devices were implanted into the animals at gestational day 13.5-14.5 and the animals were allowed to recover for 3 days. The animals were then recorded continuously during parturition. (C) HoxB8^Cre^ PIEZO1/2 knockout animals have prolonged labor, and some could not complete delivery at day 20. (D) The average interval between pup delivery was significantly longer for knockout animals in t-test. For animals that did not complete labor, one additional pup was assumed to be delivered at the end (noon of GD20). (E) Initiation of labor as marked by appearance of the first pup was delayed in knockout animals. (F) Fisher’s exact test suggests an association between PIEZO knockout and delayed labor initiation. (G) Example intrauterine traces indicate that contractions from the knockout animals are weaker during delivery. (H) Contractions in the 2-hour window after the appearance of the first pup are shown in the heatmap. (I) Fewer strong contraction peaks were observed for all knockout animals. (J) The average pressure for contraction peaks is significantly lower in knockout animals using t-test. Data quantified from 9 animals in each group.

We recorded parturitions from *Piezo1* and *Piezo2* double knockout animals and their control littermates. The labor process in knockout animals was slower, as most control animals completed all deliveries within 2 hours, but the knockout animals delivered over an extended period of time (Fig. 3C). Consequently, the average interval between pups for the knockout animals was significantly longer than control (Fig. 3D), consistent with prolonged labor. The knockout animals also tended to delay the delivery of their first pup, with approximately half delivering after zeitgeber time 3 (ZT3), whereas all control animals initiated delivery before ZT3 (Fig. 3, E and F).

The intrauterine pressure measurements further suggested that the knockout animals had weaker uterine contractions during delivery. Before the onset of parturition intrauterine pressure in both knockout and control animals was overall low with occasional peaks in intrauterine pressure, but after the delivery of the first pup, control animals had frequent high-pressure peaks indicative of strong uterine contractions (Fig. 3G; Fig. S3, A and B). When zoomed into the 2-hour time window after the appearance of the first pup, contractions in the knockout animals were minimal in contrast to frequent contractions in the control (Fig. 3H). The average pressure of the peaks from the knockout animals was significantly lower compared to the control (Fig. 3, I and J). Nonetheless, most of the knockout animals eventually exhibited compensatory increases in contraction strength (Fig. S3, C and D), suggesting that other mechanisms may still activate strong uterine contractions in the absence of PIEZO1/2. Taken together, these observations suggest that PIEZO channels are critical for robust uterine contractions during labor.

### *Piezo1/2* in the uterus and sensory nerves function complementarily

Because *Piezo1* is primarily expressed in reproductive tissues and *Piezo2* in sensory neurons, their apparent compensatory interactions likely arise from distinct mechanisms in these separate compartments. To pinpoint the specific contributions of PIEZO channels in the uterus and sensory neurons, we employed a combinatorial knockout strategy. We generated uterine knockouts using PGR^Cre^, which efficiently targets the reproductive tract (*10*), and confirmed knockout efficiency by smFISH (Fig. S4, A to C). Since Piezo2 is required for respiration, existing sensory neuron-specific Cre lines either cause lethality or only target a limited subset of DRG neurons. To overcome these limitations, we delivered AAV expressing Cre intrathecally to target the caudal DRG below the T13 level, a range closely matching that targeted by HoxB8^Cre^, thus preserving respiratory function while broadly targeting sensory neuron subtypes.

This approach enabled us to generate animals with uterus-specific, DRG-specific, and combined knockouts, along with corresponding control groups (Fig. 4, A and B). To avoid complications during mating and early gestation, virus was delivered after embryo implantation (GD 5.5–6.5). Around GD 14.5, the pregnant animals receiving Cre virus began to exhibit signs of proprioceptive defects, consistent with the expected phenotype resulting from PIEZO2 deletion in proprioceptive sensory neurons.

**Figure 4.**
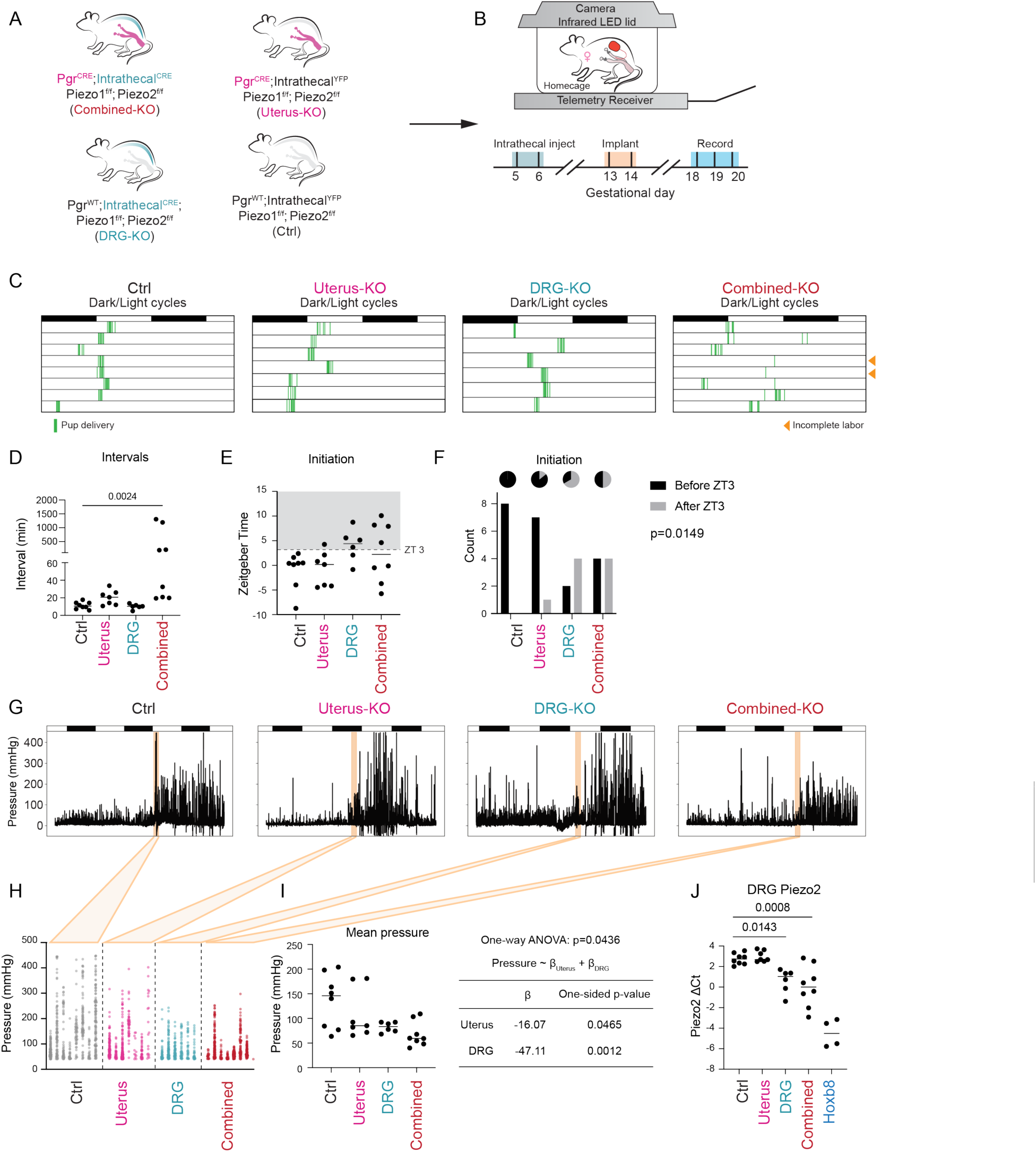
PIEZO in both the uterus and the sensory neurons contribute to labor defects. (A) PGR^cre^ is used to target the reproductive tract while MacPNS1 AAV encoding Cre is delivered intrathecally to target the sensory neurons. PGR^WT^ animals and AAV encoding YFP were used for corresponding controls. (B) The AAV injection is given at gestational day 5.5-6.5 to avoid affecting mating and pre-implantation events. The telemetry implants are placed at day 13.5-14.5. After 3 days of recovery, the animals are placed under recording for parturition. (C) The animals with PIEZO1/2 deletion in both the uterus and the DRGs have prolonged labor, while deletion in the uterus or DRG alone has less severe effect. (D) The mean intervals between pups are significantly increased in combined knockout animals (one-way Brown-Forsythe ANOVA test). Uterus-specific knockouts also exhibited a trend towards longer intervals while DRG alone deletion is almost the same as the control. For animals that did not complete labor, one additional pup was assumed to be delivered at the end (noon of GD20). (E) The initiation of labor marked by the appearance of the first pup was delayed in DRG-specific and combined knockouts. (F) Fisher’s exact test indicates that initiation after Zeitgeber time 3 (ZT3) is significantly associated with DRG-specific or combined knockouts. (G) Example intrauterine pressure traces suggest a trend of decreased contractions for knockout animals. (H) The pressure peaks within 2-hour window of delivery for each animal were curated into the plot. (I) The mean pressure for the peaks of each animal was significantly different across groups (one-way Brown-Forsythe ANOVA test). The effect size from uterus-specific and DRG-specific knockouts was then estimated using generalized least square (GLS) model. (J) Piezo2 expression in L6 and S1 level DRGs was analyzed by qPCR. Animals receiving Cre viruses have significantly lower Piezo2 expression (one-way Brown-Forsythe ANOVA test), but still higher than HoxB8^Cre^ mediated knockouts.

The labor progression differed significantly between treatment groups (Fig. 4C). Ablation of *Piezo1/2* in the uterus or DRG alone had limited effect on the overall process, whereas combined deletion in both tissues resulted in prolonged labor, resembling the phenotype observed in HoxB8^Cre^-mediated knockout. Two animals in the combined knockout group even failed to complete labor by GD 20. The average interval between pup deliveries was significantly longer in combined knockouts, whereas DRG-specific knockouts exhibited intervals similar to controls (Fig. 4D). Uterus-specific knockouts also showed a trend towards prolonged labor. Furthermore, labor initiation, marked by the appearance of the first pup, was delayed in both DRG and combined knockouts (Fig. 4, E and F). These phenotypic changes closely mirrored those observed for HoxB8^Cre^-mediated knockout animals, albeit to a lesser extent. Overall, the roles of *Piezo1/2* in DRG neurons and the uterus appear to be additive, with combined deletion leading to more pronounced labor defects.

Intrauterine pressure measurements further supported an additive effect of PIEZO channels from the uterus and DRG neurons (Fig. 4G). Within the 2-hour window following labor onset uterine contractions were progressively weaker from controls to uterus-specific, DRG-specific, and finally combined knockouts (Fig. 4, H and I). When analyzed using a generalized least squares (GLS) model treating uterus-specific and DRG-specific knockouts as two independent factors, deletion of PIEZO channels in either compartment significantly reduced contraction intensity. The estimated effect size was -16.07 mmHg for uterine knockout and -47.11 mmHg for DRG knockout. Notably, this effect was specific to the 2-hour window after labor onset, as contraction intensity prior to labor (Fig. S4, C), or beyond 2 hours after the first pup (Fig. S4, D) remained comparable across groups. Moreover, although *Piezo2* expression in caudal DRG (L6 and S1 levels) was significantly reduced in AAV-mediated knockout animals, its level varied between animals and remained higher than in HoxB8^Cre^ knockout animals (Fig. 4J), consistent with the milder phenotypes observed in AAV-mediated models. Taken together, our data demonstrate that PIEZO channels in both the reproductive tract and DRG neurons contribute additively to normal labor progression and uterine contractility.

### *PIEZOs are* required for upregulation of gap junctions in the uterus

PIEZO1 and PIEZO2 function as low threshold mechanosensors that mediate Ca^2+^ influx. Notably, uterine contractions occurring as early as one day before labor generate pressures exceeding the half-maximal activation threshold (P50) of PIEZO1 (−28.0 ± 1.8 mm Hg (*30*)), suggesting that these channels are likely activated well before labor onset. Such early activation is poised to not only transiently alter intracellular calcium dynamics but also initiate longer-term transcriptional changes. To test this hypothesis and identify PIEZO-dependent transcriptional changes in the uterus, we performed single-nuclei RNA sequencing (snRNA-seq) on uterine horns from GD18.5 knockout and control animals (*HoxB8*; P1; P2 or control littermates) (Fig. 5A; Fig. S5A).

**Figure 5.**
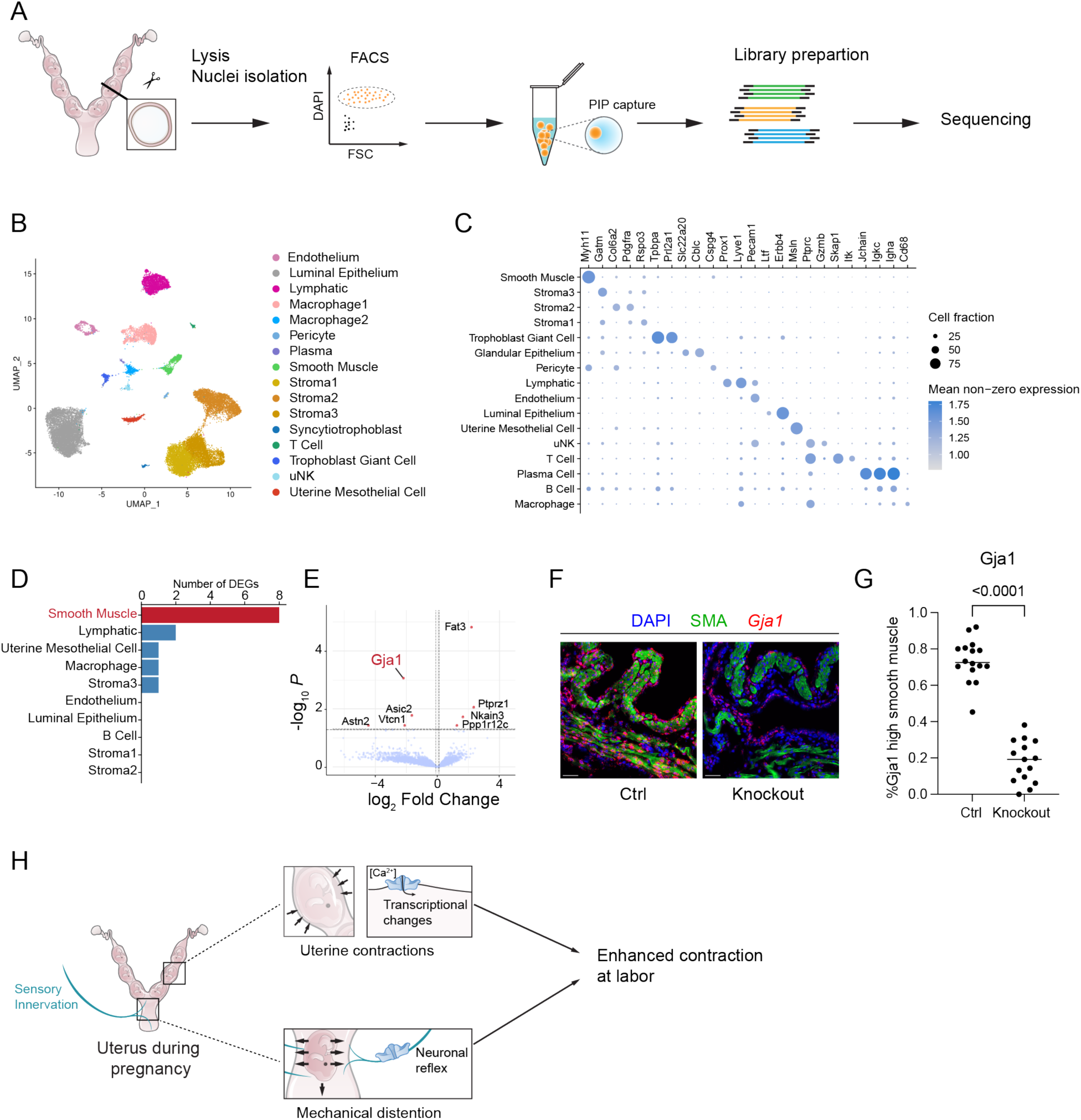
Single nuclei sequencing of the uterus revealed a defect in gap junction upregulation. (A) Uterus samples at gestational day 18.5 were dissected out, isolated from the placenta and the fetus, and processed for single nuclei sequencing. (B) Major cell types of the uterus are identified from sequencing, based on the known markers in (C). (D) A limited number of differentially expressed genes are identified between major cell types of the control and *Piezo1/2* knockout samples. Most DEGs are in the smooth muscle cells. (E) The DEGs in smooth muscle is shown with fold change and adjusted p-value. Gap junction *Gja1* is expressed at lower level in the knockout samples. (F) smFISH of *Gja1* combined with immunofluorescence of smooth muscle actin (SMA) revealed that the *Gja1* expression is dampened in knockout samples as quantified in (G) using t-test. Scale bar: 50 µm. (H) The overall role of *Piezo1/2* in parturition. *Piezo1/2* in the uterus mediates calcium influx, leading to transcriptional changes that enhance contractility, while sensory innervation mediates neuronal reflexes that promote uterine contraction. The two compartments cooperatively promote parturition.

After clustering nuclei based on their transcriptomes, we visualized the data in a 2D plot generated by Uniform Manifold Approximation and Projection (UMAP) (Fig. 5B; Fig. S5B). Major uterine cell types were well separated in the plots and expressed their corresponding marker genes (Fig. 5C). Nuclei from different animals or different genotypes showed substantial overlap in the UMAP plots (Fig. S5, C and D), suggesting that *Piezo1/2* deletion did not disrupt uterine development. This observation aligns with our earlier findings that knockout animals experience normal gestation and is further supported by cell proportion analysis (Fig. S5E), which revealed similar fractions of all cell types in control and knockout uterus samples.

We next investigated differentially expressed genes (DEGs) in each cell type. As expected for PIEZOs as ion channels that influence transcription indirectly through cumulative cation influx, only a small number of DEGs were identified (Fig. 5D).

Interestingly, most of the DEGs were detected in smooth muscle cells, the primary mediators of uterine contractions in labor. These genes may be regulated by intracellular Ca^2+^ dynamics and are thus sensitive to the loss of PIEZO activity. Among the DEGs, gap junction protein alpha1 (*Gja1*, also known as connexin 43), which is specifically expressed at late gestation (*31*), was significantly reduced in smooth muscles cells from knockout animals (Fig. 5E). Notably, previous studies have shown that smooth muscle-specific *Gja1* knockout mice exhibit delayed, and occasionally incomplete labor (*31*), aligning with our observations in *Piezo1/2* double knockout animals. We further validated the reduction of *Gja1* expression in GD 18.5 uteri from knockout animals using smFISH. Although Gja1 was minimally expressed in non-pregnant uteri (Fig. S5), its expression increased on GD 18.5, with control animals displaying significantly higher levels in smooth muscle cells (Fig. 5, F and G). These findings suggest that PIEZO channels play a critical role in promoting *Gja1* upregulation before labor, and that reduced *Gja1* expression contributes to impaired coordination in uterine contractions in the PIEZO knockouts.

## Conclusion

Our findings provide molecular evidence that mechanotransduction plays a vital role in mammalian parturition. PIEZO1 and PIEZO2 are critical to this process, operating with multiple layers of functional redundancy. At the gene level, each channel can compensate for the loss of the other. At the organ systems level, PIEZO channels in the reproductive tract and sensory neurons carry out distinct but complementary roles (Fig. 5H). Collectively, these mechanosensitive channels translate mechanical signals into coordinated biological responses necessary for the timely and effective progression of labor. This function is likely mediated in part through PIEZO-dependent regulation of intracellular calcium dynamics in smooth muscle cells, which in turn influences the expression of key contractility-associated genes such as *Gja1*.

## Discussion

PIEZO1 and PIEZO2 have been established as the main mechanosensitive channels in multiple cell types. In smooth muscle cells, PIEZO1 acts as the primary mechanosensitive receptor that mediates blood pressure-dependent remodeling, and genetic deletion of PIEZO1 abolishes stretch-activated currents completely (*32*). Similarly, PIEZO2 functions in sensory neurons to detect tactile pressure (*33*), bladder volume (*34*), gastrointestinal stretch (*35*) and other mechanical stimuli, and its ablation silences mechanical responses in corresponding neurons (*34–36*).

Building on these foundational roles, our study explored a novel function of PIEZO channels in reproductive physiology, revealing their regulation of parturition through overlapping, yet distinct, mechanosensory pathways. We found that while individual deletion of PIEZO1 or PIEZO2 produced only mild effects, simultaneous knockout of both channels resulted in severe defects, suggesting that PIEZO1 and PIEZO2 have complementary functions, where each can compensate for the other to some extent during parturition.

Mechanotransduction during parturition operates in at least two key domains: the reproductive tract and its sensory innervation. In the reproductive tract, uterine contractions generate mechanical forces that activate PIEZO channels, a process that may begin well before labor onset. Consistent with a role in early mechanical sensing, we observed PIEZO-dependent upregulation of *Gja1*, which encodes connexin 43, a protein crucial for coordinating uterine contractions. Previous studies have shown that transcription of *Gja1* is induced by mechanical stretch but suppressed by progesterone signaling (*37*). Moreover, intracellular calcium regulates transcription by modulating phospholipase C (PLC), NF-κB, and other signaling mechanisms, many of which have been reported to influence gene expression in the myometrium (*38, 39*). Thus, calcium influx through PIEZOs may facilitate translating mechanical stretch into transcriptional regulation of contractility-associated genes. In addition, the persistence of *Gja1* transcripts—albeit at significantly reduced levels—in the GD18.5 myometrium of *Piezo1/2* knockouts suggests that PIEZO-independent mechanism can still partially activate expression of contractility-associated genes.

In parallel, sensory innervation of the reproductive tract, especially in the vaginal region, constitutes another mechanosensory mechanism during parturition. During labor, mechanical distension of cervical and vaginal walls due to fetal passage likely activates sensory neurons via PIEZO2. This activation may initiate a neuronal reflex known as the Ferguson reflex to enhance uterine contractions and facilitate labor.

Previous studies have demonstrated that vaginal mechanical stimulation promotes uterine contraction in multiple mammalian species including humans (*12, 40–46*). Given that PIEZO2 is essential for mechanosensation in sensory neurons across various contexts (*29, 33–35*), it is well positioned to serve as the key sensor that mediates this reflex.

Our study also underscores the extensive complexity of mechanotransduction during parturition. While our experiments using PGR^Cre^-mediated uterine knockout combined with AAV-mediated DRG knockout produced parturition phenotypes similar to those seen in HoxB8^Cre^ knockouts, the latter exhibited more severe labor defects. This difference may be partially attributed to lower knockout efficiency achieved with viral delivery, but it is also possible that additional cell types targeted by HoxB8^Cre^ were not affected by our combined knockout strategy.

Additional complexity also exists within the reproductive tract and sensory innervation, the two primary compartments we focused on. Within the reproductive tract, our models did not target all cell types, such as endothelial cells (both vascular and lymphatic), which are critical for gestation (*47*) and known to engage in PIEZO-dependent signaling (*5*). Similarly, sensory innervation includes many calcitonin gene-related peptide (CGRP)-subtype sensory fibers, known to be polymodal and capable of integrating sensory inputs from multiple receptors (*48*). A subset of CGRP neurons can respond to high-threshold mechanical stimuli independently of PIEZO2 (*48*), suggesting the presence of alternative mechanosensitive pathways.

Moreover, loss of PIEZO1 and PIEZO2 did not eliminate all mechanical force-associated changes during pregnancy, including uterine hypertrophy and cervical remodeling. These observations suggest that additional mechanosensitive pathways contribute to the regulation of pregnancy and parturition, especially during early pregnancy. Indeed, TRP channels (*18, 20*), Hippo/YAP signaling (*16*), MAPK signaling (*14*) and several other pathways can also respond to mechanical forces and mediate key adaptations in pregnancy. Building a comprehensive network that integrate these mechanosensitive pathways with PIEZO-mediated processes will be a critical direction for future research.

While our study demonstrates that PIEZO channels are crucial for parturition in mice, direct validation of our findings in humans remains challenging due to severe physiological consequences associated with PIEZO loss-of-function mutations. PIEZO1 knockout mice exhibit embryonic lethality (*49*), and PIEZO2 deficiency in humans results in profound deficits in proprioception, tactile sensation and interoception (*50*).

However, accumulating evidence suggests that mechanotransduction pathways are conserved across species. PIEZO1 and PIEZO2 exhibit conserved expression patterns in both mouse and human myometrium, and GJA1 remains the predominant connexin protein expressed in human myometrium (*51*). Moreover, calcium signaling is universally central to uterine contractions across mammalian species. Clinically, nifedipine, a commonly used tocolytic drug, exerts its effect by inhibiting voltage-gated L-type calcium channels (*52*). Given the conserved expression patterns of PIEZO1 and PIEZO2 in mouse and human myometrium, along with the universal importance of calcium signaling in uterine contractions, it is plausible that PIEZO channels function cooperatively with L-type calcium channels, raising the intriguing possibility that pharmacological inhibition of PIEZO channels could enhance the efficacy of existing tocolytic therapies. Physical or pharmacological activation of PIEZO might in turn facilitate labor induction. Further studies are essential to explore this potential therapeutic avenue.

## Acknowledgments

We thank all members of the Patapoutian lab for their support and feedback, Jeanine Ahmed and Sebastian Burquez for assisting with experiments. We thank Robert Froemke and Luisa Schuster for valuable advice on video recording of mice behavior. We thank Nirao Shah for sharing the PGR-cre mouse strain. We thank Lindsay Burnett and Mala Mahendroo for insightful suggestions during the project. Microscopy was performed in the Scripps Research Core Microscopy Facility and the Nikon Center of Excellence at Scripps Research. We also acknowledge the staff at Scripps FACS core and genomics core for sample preparation. This study was supported in part by the WashU Reproductive Specimen Processing and Banking Biorepository (ReProBank).

## Materials and Methods

### Animals

Mice were group-housed in standard housing with 12:12h light:dark (light on at 6 am, off at 6 pm) with ad libitum access to chow diet and water unless specified. Light-on time is defined as zeitgeber time (ZT) 0. The room temperature was kept around 22 °C and humidity between 30-80% (not controlled). Mice for telemetry recordings were single housed. Mice 8-12 weeks of age from the following strains were used for this study: wild-type (WT) C57BL/6J (Jackson #000664), PIEZO2-Cre (Piezo2^tm1.1(cre)Apat^, Jackson #027719)(*53*), HoxB8-Cre (MGI:4881836)(*54*), PGR-Cre (Jackson Pgr^tm1.1(cre)Shah^/AndJ, Jackson #017915)(*55*), PIEZO2^fl/fl^ (Piezo2^tm2.2Apat^/J, Jackson #027720)(*56*) and PIEZO1^fl/fl^ (Piezo1^tm2.1Apat^/J, Jackson #029213)(*57*). All animals were maintained on C57BL/6J background, except for PIEZO2-Cre, which were on CD-1;C57BL/6J background. Only female animals were used for pregnancy-related studies. During mating, each female mice was housed with a WT male mice after 4 pm (ZT10).

Pregnancy was confirmed by identification of semen plug before 8 am (ZT2) and that morning was recorded as gestational day 0.5 (GD0.5). All animal use protocols were approved by The Scripps Research Institute Institutional Animal Care and Use Committee and were in accordance with the guidelines from the NIH.

### Human Myometrial Samples

De-identified human myometrial tissues were obtained from the Reproductive Specimen Processing and Banking Biorepository at Washington University in St. Louis. Inclusion criteria for samples used in this study were gestational age ≥37 weeks, singleton gestation, and nulliparity. Additional exclusion criteria were known HIV, hepatitis B or hepatitis C infection. Human myometrial biopsies were collected from the lower uterine segment of the hysterotomy performed at the time of scheduled cesarian delivery, prior to labor. Samples were transported on ice, flash frozen in liquid nitrogen within 60 minutes of collection and stored at -80 °C until further analyses. Use of human samples was under the approval of the Scripps Research Institute Institutional Review Boards (IRB-23-8255).

### Adeno-Associated Viruses (AAVs)

MacPNS.1-CAG-DIO-mScarlet (capsid Addgene #185136(*58*)) were used to label Piezo2+ sensory neurons. MacPNS.1-CAG-iCre-YFP, MacPNS.1-CAG-YFP were used to generate DRG-specific knockouts and control groups for functional studies. The AAV cargo vectors were cloned, and packaged into AAV in-house following a published protocol. After titration by qPCR, AAV was aliquoted into 10 μL and flash frozen for long-term storage.

### Surgeries

Isoflurane (4% for induction, 1.5-2% for maintenance) was used for general anesthesia. The surgery region was shaved, cleaned and sterilized with ethanol and povidone-iodine. Post-operatively, mice received a subcutaneous injection of flunixin and topical antibiotic ointment for pain management and infection prevention.

The telemetry device used was HD-X11 from Data Science International (DSI) and the surgery followed previously published procedure(*59*). To implant telemetry devices, an incision along the midline was made to each animal at GD13.5-14.5 and the uterus was gently retracted. At least 3 embryos in gestation need to be present in one side of the uterine horn for implantation. A small incision just enough for the entry of the pressure catheter was made in the uterine horn at a region between embryos and the pressure catheter from the device was inserted past 1.5 embryos. The catheter was then secured in place by adding 1 µL of tissue adhesive at the site of entry. If necessary, the uterus was close with one suture using 8-0 nylon suture. The abdominal incision was closed with suture afterwards. For prophylaxis of infection, sulfamethoxazole and trimethoprim was added to the drinking water one day before and two days after the surgery.

For CTB retrograde labeling, each animal received 4–5 μL of 0.1% CTB-488 (Invitrogen C34775) or CTB-647 (Invitrogen C34778) in PBS. The injections were made with glass pipettes mounted onto Hamilton syringes and delivered 0.2-0.5 µL per site. Multiple injections were made throughout the entire uterus/vagina. DRGs were harvested 3-4 days post-surgery for histology.

For intrathecal injection, 8-10 µL of AAV was delivered at L5-L6 junction using a Hamilton syringe. For sensory nerve labeling, MacPNS.1 CAG-DIO-mScarlet AAV was injected into PIEZO2-Cre mice at a total dose of 6E11 vg. The tissues were collected 3- 4 weeks after injection. For DRG-specific knockouts in functional assays, MacPNS.1 CAG-YFP or MacPNS.1 CAG-iCre-YFP was delivered into animals at GD5.5-6.5 with a total dose of 3E11 vg.

### Preparation of single nuclei sequencing

Animals were sacrificed at 10-11 am on GD18.5 under deep anesthesia with isoflurane and the uterus was dissected out and placed in ice-cold PBS. The fetus along with placenta and amniotic membranes was gently peeled off and the uterus was tapped dry on Kimi wipe before frozen in liquid nitrogen. On the day of sequencing sample preparation, the frozen samples were pulverized with a pestle chilled with liquid nitrogen. All buffers used in later steps contain 0.5 U/ml RNase inhibitor (3335399001, Sigma-Aldrich). The tissues were then resuspended in lysis buffer (21mM MgCl2, 1mM CaCl2, 146mM NaCl, 10mM Tris pH 8, 0.02% Tween-20, 1% Polyvinylpyrrolidone) and lysed in a pre-chilled dounce homogenizer. Homogenized tissue was passed through 70 µm cell strainer to remove large debris and then spun down at 500 g for 5 min.

Nuclei was resuspended in wash buffer (1mM MgCl2, 1mM CaCl2, 146mM NaCl, 10mM Tris pH 8, 1% Polyvinylpyrrolidone) containing 5 µg/ml DAPI before loading onto Sony MA900 Sorter equipped with a 100-µm microfluidic sorting chip. Single nuclei were sorted into a collection tube and then centrifuged at 500 g, 5min at 4 C. After resuspension in NSB (Fluent BioSciences) and counting on a hemacytometer slide, 5000 nuclei were taken for PIPseq library preparation using PIPseq T2 v4.0PLUS 3’ kit (Fluent BioSciences) following the manufacturer’s instructions (*60*). The library was sequenced on Element Aviti sequencer following the Adept Workflow.

### Tissue clearing and staining

Tissues were cleared for imaging following the HYBRiD clearing protocol(*61*). In brief, the reproductive tract was dissected out after trans-cardial perfusion of ice-cold PBS and then 4% PFA (Electron Microscopy Sciences, 15714). The samples were then subjected to organic clearing by THF/25% Quadrol gradient and DCM. After embedding into A1P4 hydrogel (1% acrylamide, 0.125% Bis, 4% PFA, 0.025% VA-044 (w/v), in 1x PBS), the samples were then cleared with LiOH-Boric-SDS buffer until translucent.

The tissues were washed extensively in PBS before immunolabeling. The tissues were incubated in Rabbit anti-RFP antibody (Rockland #600-401-379, 1:500) in PBST (1 x PBS, 0.1% Tween-20, 0.1% Triton X-100) for 5 days at RT and then washed with PBST. This was followed by incubation with Donkey anti-Rabbit-AlexaFlour PLUS 647 antibody (ThermoFisher, A32795, 1:1000) for 5 days at RT, and sequential washed in PBST. The samples were then equilibrated in EasyIndex (RI 1.52, LifeCanvas) for confocal and light sheet microscopy imaging.

### RNA Fluorescence in Situ Hybridization (RNAscope)

Mice were deeply anesthetized by isoflurane prior to perfusion with 20-25 ml ice-cold PBS. Tissues were dissected and then fixed in ice-cold 4% PFA 1-2 hr, before transferring to 30% sucrose for dehydration overnight. The tissues were then embedded in Tissue-Tek OCT Compound (Sakura Finetek). The uterus samples were cryosectioned at 15 μm and collected onto SuperFrost Plus Microscope Slides (Fisher Scientific 12-550-15). The target transcripts were detected with RNAscope Multiplex Fluorescent Reagent Kit v2 Assay (Advanced Cell Diagnostics) following the manufacturer’s protocol for fixed-frozen tissues. Human tissue samples were directly embedded for cryosection and then stained following the manufacturer’s protocol for fresh frozen tissues. The following probes were used: Mm-Piezo2-E43-E45 (Advanced Cell Diagnostics 439971), Mm-Piezo1-O1 (500511, Advanced Cell Diagnostics), Mm- Gja1 (Advanced Cell Diagnostics 486191) for mouse tissues, and Hs-PIEZO1 (Advanced Cell Diagnostics 485101), Hs-PIEZO2 (Advanced Cell Diagnostics 449951) for human tissues. Cell markers were stained with antibody afterwards for smooth muscle actin (Acta2, Abcam ab267536) or Lyve1 (R&D systems AF2125). Sections were mounted using Prolong Glass Antifade Reagent (ThermoFisher P36982) and imaged under a 20X objective on a Nikon AX confocal microscope.

### Tissue histology

Mice were deeply anesthetized by isoflurane prior to perfusion with 20-25 ml ice-cold PBS. Tissues were dissected and then fixed in ice-cold 4% PFA overnight and then dehydrated in 70% ethanol. The tissues were then sent for paraffin embedding, sectioning and staining at the Sanford Burnham Prebys Histology Core. The stained slides were imaged with a Leica Asperio AT2 slide scanner.

### Confocal microscopy

Mounted uterus samples were imaged with Olympus FV3000 confocal microscope using the following objectives: 4X, 0.28 NA, air (XLFluor, Olympus); 10X, 0.6 NA, water immersion (XLUMPlanFI, Olympus). Images were acquired with Fluoview (v2.4.1.198).

RNAscope samples and DRG samples were imaged with Nikon AX confocal microscope with a 20x objective or a 16x water immersion objective.

### Light sheet microscopy

The uterus samples were mounted using 1% agarose/EasyIndex. The samples were equilibrated in SmartSPIM chamber filled with EasyIndex Matched Immersion Oil (LifeCanvas) overnight before imaging. Images were acquired using a 3.6X, 0.2 NA objective (LifeCanvas) with bilateral illumination along the central plane of symmetry within the sample. The voxel size after stitching was 1.8/1.8/2 µm.

### Mouse imaging

Mice we imaged using a SainSmart 5MP 1080p Night Vision camera module with a 160 field of view (FOV) fisheye lens to capture a large field of view. Two infrared red lights were installed on both sides of the camera to enable imaging in the dark. Raspberry Pi Zero W was used to control the camera and store the recorded videos. MotioneyeOS was used to remotely control the recordings and minimize disturbance to the animals.

The camera and Raspberry Pi were housed in 3D printed capsules made with Nylon X using a Fusion 3, F410 high-performance 3D printer. The camera and Raspberry Pi capsules were printed using ESD-safe PLA 3D printing filament (3DXSTAT) to prevent electrostatic discharges (ESD).

### RNA extraction

DRG were dissected out and flash frozen in liquid nitrogen. Total RNA was extracted from frozen tissue using TRIzol (ThermoFisher 15596018) and RNeasy Micro Kits (Qiagen 74004) following manufacturer’s instructions.

### Reverse Transcription-PCR

For RT-PCR analysis, total RNA was reverse-transcribed with Maxima H Minus First Strand cDNA Synthesis Kit (Thermo Fisher K1652). PCR reaction was set up with SsoAdvanced Universal SYBR Green Supermix (Bio-Rad) on CFX384 real-time PCR system (Bio-Rad). Normalized mRNA expression was calculated using ΔCt method, using *Tbp* (encoding TATA-box-binding protein) mRNA as the reference gene. Statistics was performed on ΔCt. Primer sequences (forward and reverse sequence, 5’ → 3’, respectively) are *Tbp* (CCTTGTACCCTTCACCAATGAC and ACAGCCAAGATTCACGGTAGA); *Piezo2* (CAAAGTCAATGGTCGCGTGT and CAAGGCTGGCCGTCATATTC). QUANTIFICATION AND STATISTICAL ANALYSIS Transcript quantification in RNAscope: Maximum z-projection images were used for RNAscope quantification using CellProfiler(*62*). Cells were identified based on expanded regions from nuclei DAPI staining. Puncta from RNAscope signals were identified as objects and assigned to corresponding cell nuclei. Immunofluorescent signals such as SMA were quantified by intensity and used to differentiate cell types.

### snRNA-seq analysis

The sequencing results were obtained in fastq format. After transcriptome mapping using PIPseeker (Fluent BioSciences), the raw matrix was filtered with CellBender(*63*) to remove empty droplets and then filtered out nuclei with high mitochondria reads (>5%) using Seurat(*64*). Doublets were detected and filtered using DoubletFinder(*65*). Feature selection was performed with *FindVariableFeatures* within Seurat. Data from each sample was scaled and then integrated using Harmony(*66*). Uniform manifold approximation and projection (UMAP) was used for visualization after integration. Cell types were assigned based on markers. Cell proportions across different conditions were tested by Propeller(*67*) for difference. Differentially expressed genes in each cell type were identified using SVA-EdgeR following a previously published procedure(*68*).

### Telemetry analysis

Intrauterine pressure data was exported in ASCII format from Ponemah 5.20. Pup delivery was manually annotated from video. The time for birth was determined with the accuracy of 1 min. Pressure data was processed with a custom Python script, and resampled to a sampling rate of 1 per second. The pressure peaks with a minimum height of 40 (mmHg) and prominence of 20 were identified using scipy.find_peaks function and statistical tests (t-test and one-way ANOVA Brown-Forsythe test) were conducted in GraphPad Prism. To analyze the effect of uterus-specific and DRG-specific knockouts on peak pressures, average peak pressures from animals in the 4 treatment groups (control, uterus-specific, DRG-specific and combined knockouts) were considered in a linear model for the continuous outcome 𝑷_𝒊_, the average peak pressure for the 𝒊th animal, which depends on two categorical predictors 𝑫_𝑫𝑹𝑮_ and 𝑫_𝑼𝒕𝒆𝒓𝒖𝒔_ denoting whether PIEZO1/2 is deleted from DRG or uterus of this animal. For observation 𝒊, the model is

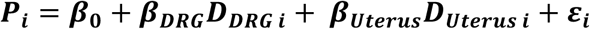

The error term 𝜺_𝒊_ is assumed to follow a normal distribution with mean 0 but with a variance that depends on the group:

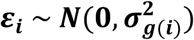

Here, 𝒈(𝒊) indicates the treatment group to which observation 𝒊 belongs. In matrix form, the model can be written as

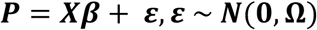

Where 𝑷 is the vector of outcomes, 𝑿 is the design matrix including the intercept and the categorical factors, 𝜷 is the vector of parameters, and 𝛀 is a diagonal covariance matrix for the error terms.

The variances are not equal across the groups (heteroscedasticity) as viral-induced knockouts are inherently more variable than Cre expressed from a genomic locus. We thus used Generalized Least Squares (GLS) to obtain efficient estimates. The GLS estimator is given by

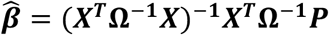

We aim to test whether the effect of predictor𝑫 (𝑫_𝑫𝑹𝑮_ and 𝑫_𝑼𝒕𝒆𝒓𝒖𝒔_) is decreasing and can be framed as a one-sided hypothesis test:

Null hypothesis: 𝑯_𝟎_: 𝜷_𝑫_ ≥ 𝟎; Alternative hypothesis: 𝑯_𝟏_: 𝜷_𝑫_ < 𝟎

The test statistic for the coefficient 𝜷_𝑫_is

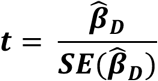

Under 𝑯_𝟎_, it follows a t-distribution with degrees of freedom approximated by 𝒏 − 𝒑 (where n is the number of observations and p is the number of estimated parameters). For a one-sided test, if the estimated coefficient β̂_𝑫_ is negative, the p-values is computed as

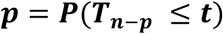

Where 𝑻_𝒏 ‒ 𝒑_ is subject to t-distribution with 𝒏 − 𝒑 degrees of freedom.

### Study design and statistics

The sample size in this study was not determined using statistical methods, but was guided by prior studies and literature in the field using similar experimental paradigms. Mice with post-surgery complications, as defined by loss of >1g body weight in 2 days after surgery, were euthanized before telemetry recording (N=2 total). One animal in HoxB8 Piezo1/2 double knockout group was excluded from analysis for lack of any signs of going into labor on GD 20.

Data collection was conducted blindly, with post hoc registration to respective condition for unbiased analysis. Animals with proprioceptive defects can still be spotted and are not truly blinded to the observer in this aspect. GraphPad Prism was used for statistical tests, unless specified otherwise. Sample sizes for each experiment are reported in the figure legends. All *in vivo* experiments were repeated at least twice or combined from at least two independent cohorts, yielding consistent results.

### Data and Code Deposition

The snRNA-seq data have been deposited at the Broad Institute Single Cell Portal under accession number SCP2924. All code used for data processing and analysis will be deposited on Github.

---

**Fig. S1.**
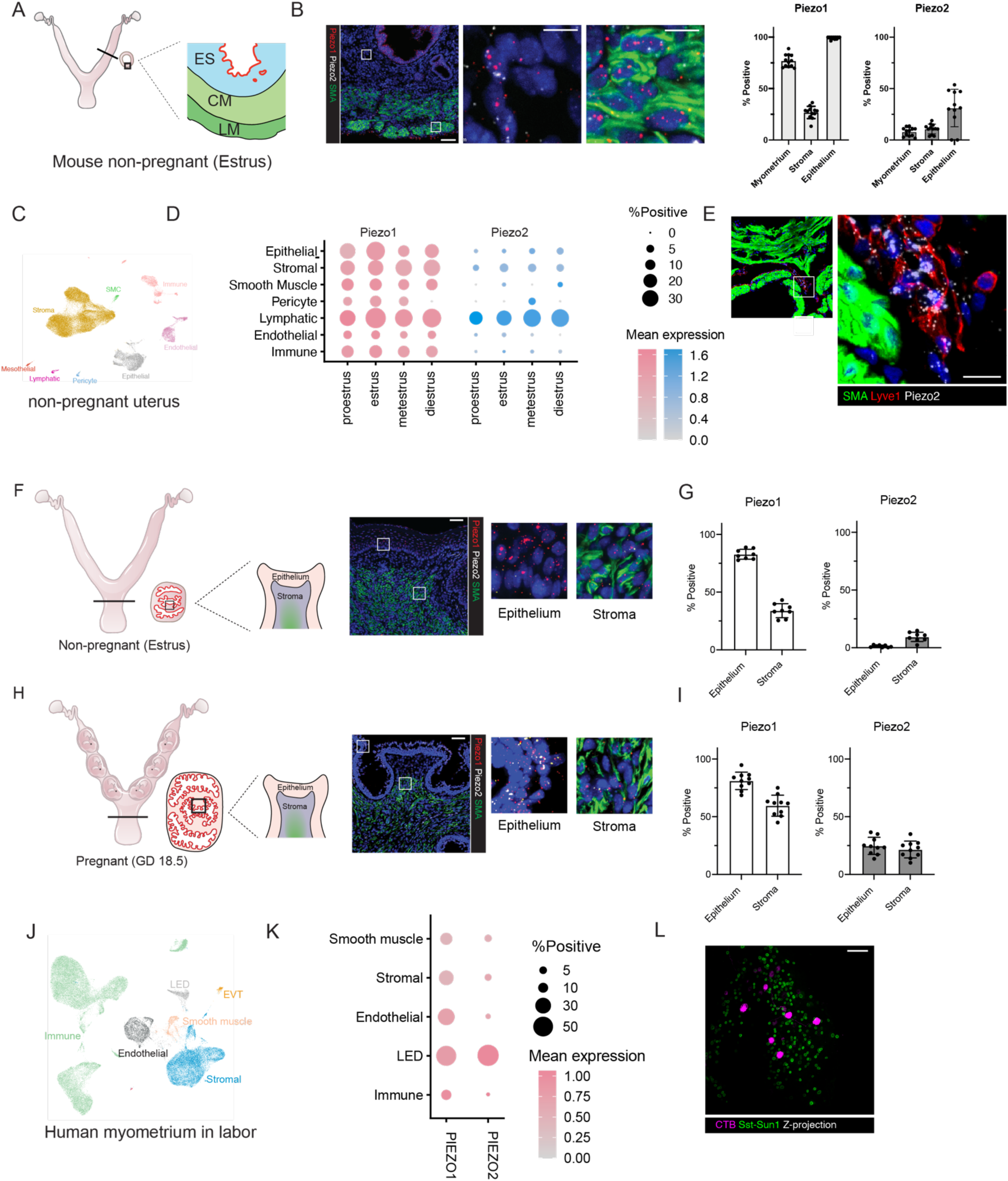
(A, B) PIEZO1/2 expression pattern in non-pregnant uterus is similar as the pregnant one. Scale bar: 50 µm in the overview, 10 µm in the inset. (C, D) Published single cell sequencing dataset reveals that Piezo1 and Piezo2 are expressed across multiple cell types and may have fluctuations according to the estrous cycle.(E) Lymphatic endothelial cells co-express PIEZO1 and PIEZO2 at high levels and are primarily localized between smooth muscle layers at GD 18.5. Scale bar: 50 µm (F, G) Piezo1 is expressed in the non-pregnant cervix, while Piezo2 is expressed at very low levels. Scale bar: 50 µm (H, I) In cervix at late pregnancy, Piezo1 expression maintains while Piezo2 expression is higher especially in the epithelium. Scale bar: 50 µm (J, K) In human myometrium in labor, PIEZO1 is expressed in all major cell types, while PIEZO2 is expressed almost only in lymphatic endothelial decidual cells (LEDs). (L) Retrograde labeling of uterus and cervix innervating DRG neurons with CTB revealed little overlap with Sst+ sensory neurons, the population that expresses Piezo1, suggesting that Piezo1 expression may be rare in sensory innervation of the reproductive tract. Scale bar: 100 µm

**Figure S2:**
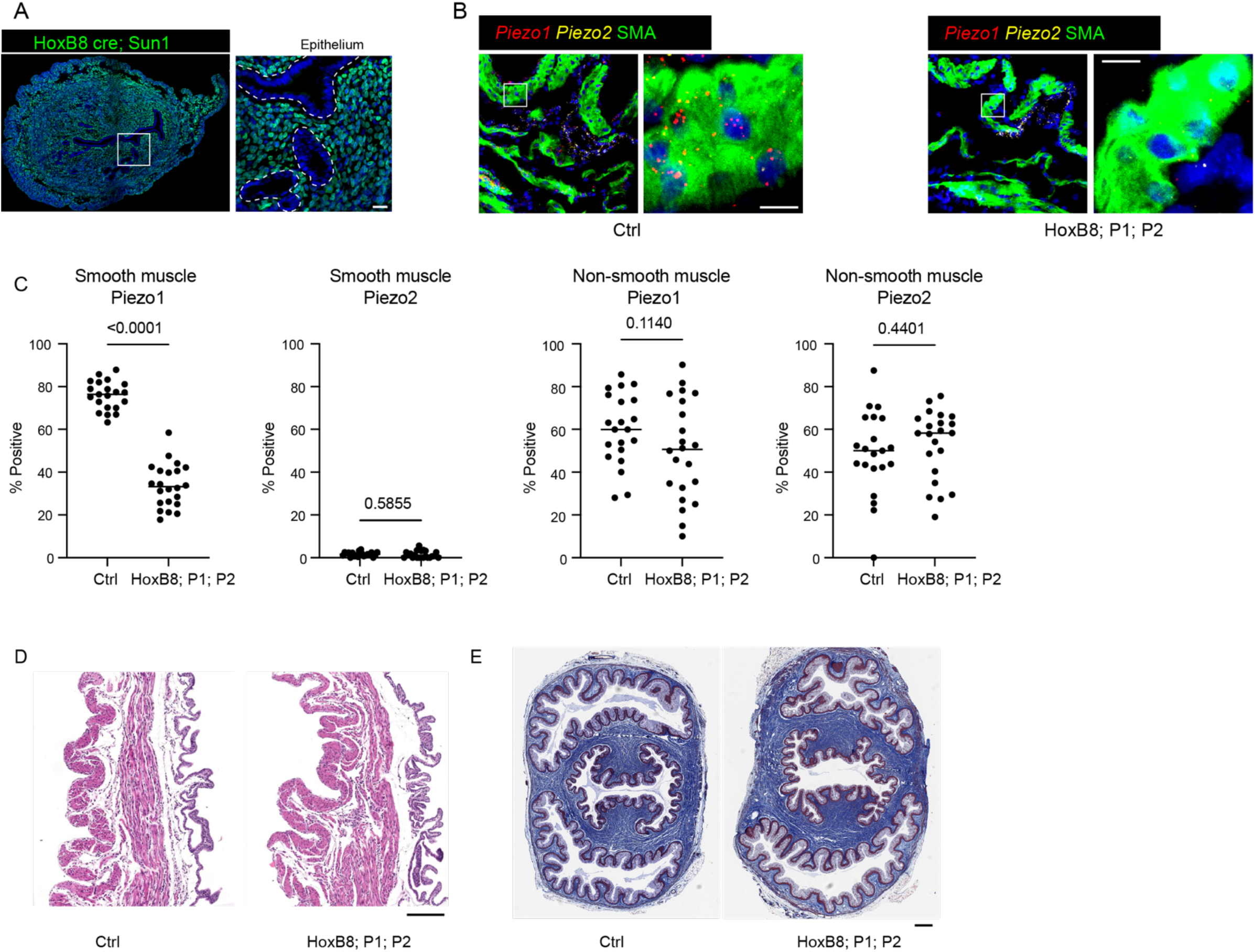
(A) HoxB8 Cre targets most cells in the non-pregnant uterus except the epithelium, as indicated by Rosa26 Sun1 reporter. Scale bar: 10 µm. (B) HoxB8 Cre effectively ablated PIEZO1/2 expression in the uterine smooth muscle cells at gestational day 18.5, but some non-smooth muscle cells that co-express PIEZO1 and PIEZO2 are not affected, as quantified in (C). Statistical test: t-test. These double positive cells are mainly lymphatic endothelial cells. Scale bar: 50 µm, 10 µm in the inset. (D) The uterus at gestational day 18.5 has similar appearance with H&E staining, suggesting that PIEZO1/2 knockout does not have a major effect on tissue structure. Scale bar: 200 µm. (E) Cervix sections at day 18.5 have similar appearance under trichrome staining, suggesting that PIEZO1/2 knockout may not affect cervix remodeling before labor. Scale bar: 200 µm.

**Figure S3.**
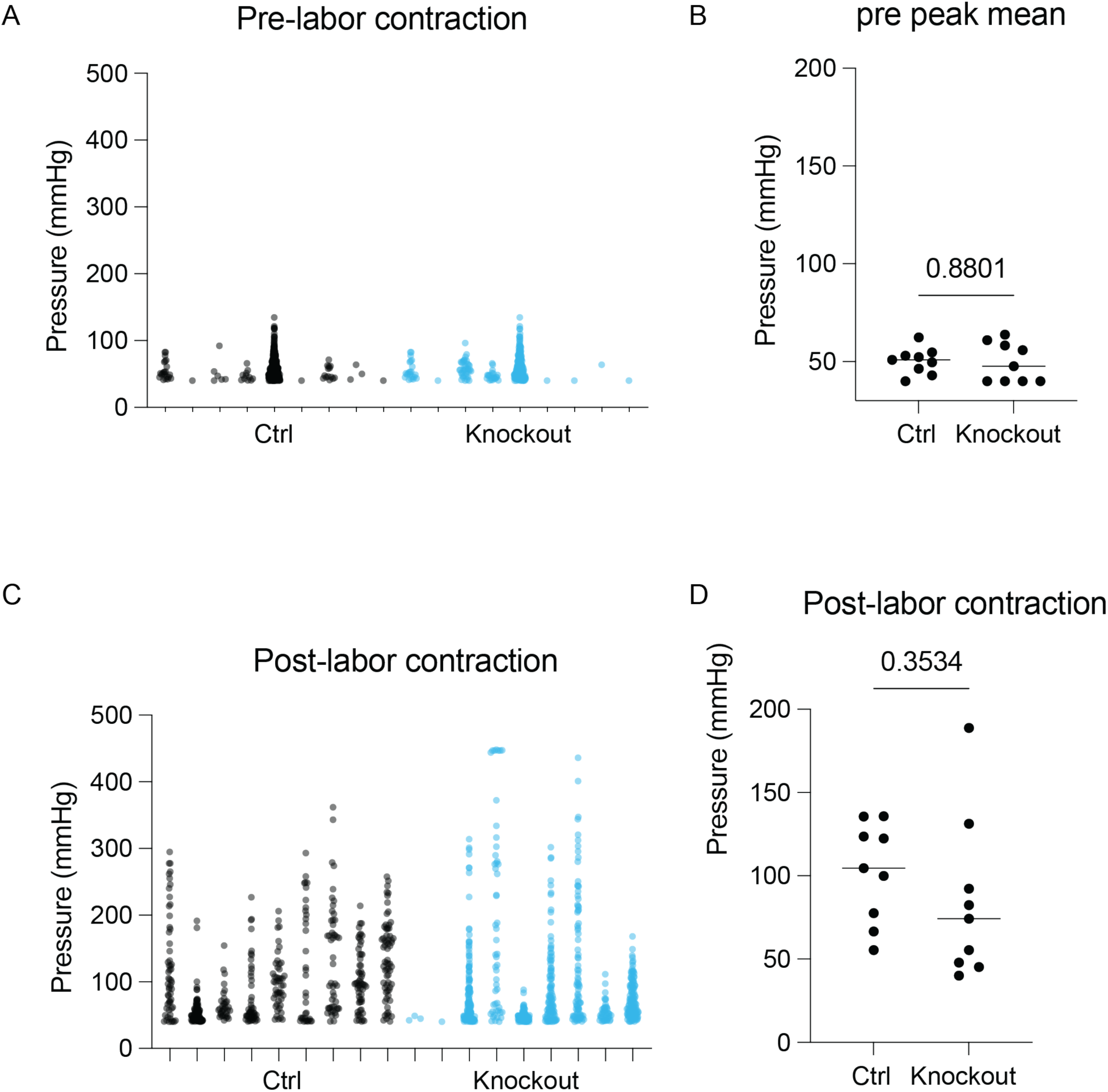
(A) Pre-labor contractions (GD 18 midnight (ZT 18) to GD18 6 pm (ZT 12)) are similar between control and knockout animals as quantified in (B). (C) Contractions 2-8hr after appearance of the first pup are similar between control and knockout animals, as quantified in D. Statistical test: t-test.

**Figure S4.**
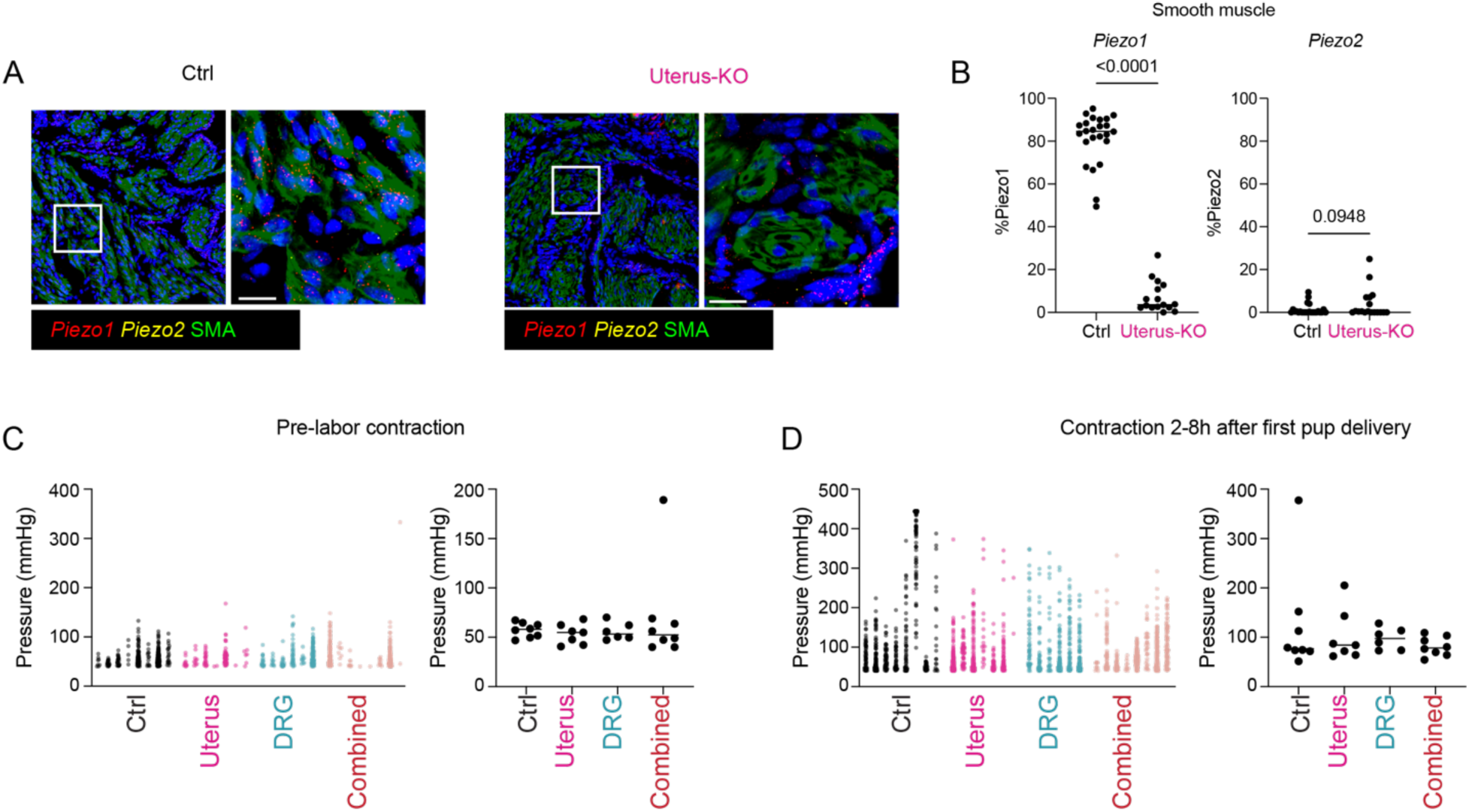
(A) Fluorescence ISH reveals that PGR cre mediates effective deletion of PIEZO1/2 in uterine smooth muscle cells, as quantified in (B). Statistical test: t-test. Scale bar: 20 µm (C) Pressure peaks from pre- labor contractions (GD 18 midnight (ZT 18) to GD18 6 pm (ZT 12)) are comparable across different treatment groups. (D) Pressure peaks at 2-8 hr after the first pup delivery are comparable across groups.

**Figure S5.**
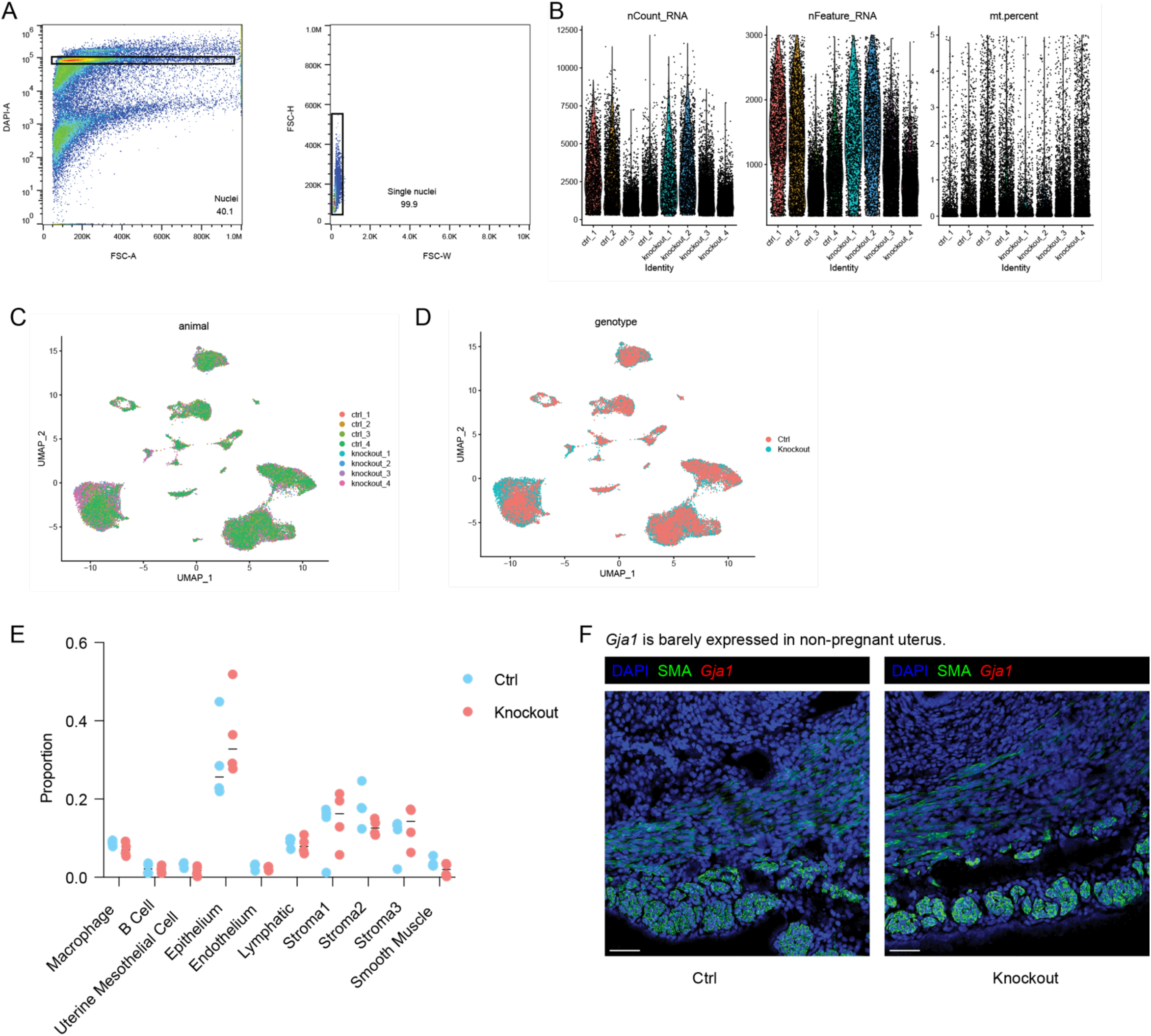
(A) The samples were gated based on DAPI in FACS to obtain nuclei and further filtered based on forward scattering light to obtain single nuclei. (B) Read depth, and sample quality metrics of the single nuclei sequencing samples are shown as violin plots. (C) Cells from different animals or different batches overlap in UMAP representation without obvious batch effect. (D) The cells from control or knockout animals intermingle in the clusters, suggesting that knockout of Piezo1/2 does not change major tissue composition, which is further confirmed by the proportions of the major cell types in (E). (E) Non-pregnant uterus from control or knockout animals had no detectable levels of Gja1, suggesting that Gja1 is upregulated at late gestation under both conditions, but the upregulation is much less in knockout animals. Scale bar: 50 µm.

## References and Notes

1. J. M. Kefauver, A. B. Ward, A. Patapoutian, Discoveries in structure and physiology of mechanically activated ion channels. Nature 587, 567–576 (2020).

2. F. Martino, A. R. Perestrelo, V. Vinarsky, S. Pagliari, G. Forte, Cellular Mechanotransduction: From Tension to Function. Front Physiol 9, 824 (2018).

3. S. Katta, M. Krieg, M. B. Goodman, Feeling force: physical and physiological principles enabling sensory mechanotransduction. Annu Rev Cell Dev Biol 31, 347–371 (2015).

4. C. C. DuFort, M. J. Paszek, V. M. Weaver, Balancing forces: architectural control of mechanotransduction. Nat Rev Mol Cell Biol 12, 308–319 (2011).

5. D. Douguet, A. Patel, A. Xu, P. M. Vanhoutte, E. Honore, Piezo Ion Channels in Cardiovascular Mechanobiology. Trends Pharmacol Sci 40, 956–970 (2019).

6. X. Di et al., Cellular mechanotransduction in health and diseases: from molecular mechanism to therapeutic targets. Signal Transduct Target Ther 8, 282 (2023).

7. B. D. Umans, S. D. Liberles, Neural Sensing of Organ Volume. Trends Neurosci 41, 911–924 (2018).

8. K. M. Myers, D. Elad, Biomechanics of the human uterus. Wiley Interdiscip Rev Syst Biol Med 9, (2017).

9. S. Jorge, S. Chang, J. J. Barzilai, P. Leppert, J. H. Segars, Mechanical signaling in reproductive tissues: mechanisms and importance. Reprod Sci 21, 1093–1107 (2014).

10. S. P. Wu, R. Li, F. J. DeMayo, Progesterone Receptor Regulation of Uterine Adaptation for Pregnancy. Trends Endocrinol Metab 29, 481–491 (2018).

11. F. J. DeMayo, J. P. Lydon, 90 YEARS OF PROGESTERONE: New insights into progesterone receptor signaling in the endometrium required for embryo implantation. J Mol Endocrinol 65, T1–T14 (2020).

12. J. K. Ferguson, A study of the motility of the intact uterus at term. Surg Gynecol Obstet 73, 359–366 (1941).

13. O. Shynlova, P. Tsui, S. Jaffer, S. J. Lye, Integration of endocrine and mechanical signals in the regulation of myometrial functions during pregnancy and labour. Eur J Obstet Gynecol Reprod Biol 144 **Suppl 1**, S2–10 (2009).

14. A. D. Oldenhof, O. P. Shynlova, M. Liu, B. L. Langille, S. J. Lye, Mitogen- activated protein kinases mediate stretch-induced c-fos mRNA expression in myometrial smooth muscle cells. Am J Physiol Cell Physiol 283, C1530–1539 (2002).

15. S. R. Sooranna et al., Mechanical stretch activates type 2 cyclooxygenase via activator protein-1 transcription factor in human myometrial cells. Mol Hum Reprod 10, 109–113 (2004).

16. K. L. Clark et al., Hippo Signaling in the Ovary: Emerging Roles in Development, Fertility, and Disease. Endocr Rev 43, 1074–1096 (2022).

17. B. Timmons, M. Akins, M. Mahendroo, Cervical remodeling during pregnancy and parturition. Trends Endocrinol Metab 21, 353–361 (2010).

18. A. Dalrymple, K. Mahn, L. Poston, E. Songu-Mize, R. M. Tribe, Mechanical stretch regulates TRPC expression and calcium entry in human myometrial smooth muscle cells. Mol Hum Reprod 13, 171–179 (2007).

19. S. D. Barnett, H. Asif, I. L. O. Buxton, Novel identification and modulation of the mechanosensitive Piezo1 channel in human myometrium. J Physiol 601, 1675–1690 (2023).

20. L. Ying et al., The transient receptor potential vanilloid 4 channel modulates uterine tone during pregnancy. Sci Transl Med 7, 319ra204 (2015).

21. P. Delmas, T. Parpaite, B. Coste, PIEZO channels and newcomers in the mammalian mechanosensitive ion channel family. Neuron 110, 2713–2727 (2022).

22. I. Winkler et al., The cycling and aging mouse female reproductive tract at single- cell resolution. Cell 187, 981–998 e925 (2024).

23. R. Pique-Regi, et al., A single-cell atlas of the myometrium in human parturition. JCI Insight 7, (2022).

24. G. Herweijer, M. Kyloh, E. A. Beckett, K. N. Dodds, N. J. Spencer, Characterization of primary afferent spinal innervation of mouse uterus. Front Neurosci 8, 202 (2014).

25. B. Xiao, Mechanisms of mechanotransduction and physiological roles of PIEZO channels. Nat Rev Mol Cell Biol 25, 886–903 (2024).

26. B. K. Tingaker, G. Ekman-Ordeberg, P. Facer, L. Irestedt, P. Anand, Influence of pregnancy and labor on the occurrence of nerve fibers expressing the capsaicin receptor TRPV1 in human corpus and cervix uteri. Reprod Biol Endocrinol 6, 8 (2008).

27. B. K. Tingaker, L. Irestedt, Changes in uterine innervation in pregnancy and during labour. Curr Opin Anaesthesiol 23, 300–303 (2010).

28. R. Z. Hill, M. C. Loud, A. E. Dubin, B. Peet, A. Patapoutian, PIEZO1 transduces mechanical itch in mice. Nature 607, 104–110 (2022).

29. R. M. Lam et al., PIEZO2 and perineal mechanosensation are essential for sexual function. Science 381, 906–910 (2023).

30. B. Coste et al., Piezo1 and Piezo2 are essential components of distinct mechanically activated cation channels. Science 330, 55–60 (2010).

31. B. Doring et al., Ablation of connexin43 in uterine smooth muscle cells of the mouse causes delayed parturition. J Cell Sci 119, 1715–1722 (2006).

32. K. Retailleau et al., Piezo1 in Smooth Muscle Cells Is Involved in Hypertension- Dependent Arterial Remodeling. Cell Rep 13, 1161–1171 (2015).

33. S. S. Ranade et al., Piezo2 is the major transducer of mechanical forces for touch sensation in mice. Nature 516, 121–125 (2014).

34. K. L. Marshall et al., PIEZO2 in sensory neurons and urothelial cells coordinates urination. Nature 588, 290–295 (2020).

35. M. R. Servin-Vences et al., PIEZO2 in somatosensory neurons controls gastrointestinal transit. Cell 186, 3386–3399 e3315 (2023).

36. A. M. Chirila et al., Mechanoreceptor signal convergence and transformation in the dorsal horn flexibly shape a diversity of outputs to the brain. Cell 185, 4541–4559 e4523 (2022).

37. C. W. Ou, A. Orsino, S. J. Lye, Expression of connexin-43 and connexin-26 in the rat myometrium during pregnancy and labor is differentially regulated by mechanical and hormonal signals. Endocrinology 138, 5398–5407 (1997).

38. C. Xu et al., Prostaglandin F2alpha regulates the expression of uterine activation proteins via multiple signalling pathways. Reproduction 149, 139–146 (2015).

39. J. A. Ingles et al., Loss of TRPV4 decreases NFkappaB-mediated myometrial inflammation and prevents preterm labor. FASEB J 39, e70418 (2025).

40. S. Sato, R. H. Hayashi, R. E. Garfield, Mechanical responses of the rat uterus, cervix, and bladder to stimulation of hypogastric and pelvic nerves in vivo. Biol Reprod 40, 209–219 (1989).

41. L. A. Clyde et al., Transection of the pelvic or vagus nerve forestalls ripening of the cervix and delays birth in rats. Biol Reprod 84, 587–594 (2011).

42. T. Higuchi, K. Uchide, K. Honda, H. Negoro, Pelvic neurectomy abolishes the fetus-expulsion reflex and induces dystocia in the rat. Exp Neurol 96, 443–455 (1987).

43. A. Klukovits, R. Gaspar, P. Santha, G. Jancso, G. Falkay, Role of capsaicin-sensitive nerve fibers in uterine contractility in the rat. Biol Reprod 70, 184–190 (2004).

44. A. P. Flint, M. L. Forsling, M. D. Mitchell, Blockade of the Ferguson reflex by lumbar epidural anaesthesia in the parturient sheep: effects on oxytocin secretion and uterine venous prostaglandin F levels. Horm Metab Res 10, 545–547 (1978).

45. J. S. Roberts, Functional integrity of the oxytocin-releasing reflex in goats: dependence on estrogen. Endocrinology 93, 1309–1314 (1973).

46. A. Shafik, Study of the uterine response to vaginal distension: the ’vagino-uterine reflex’. Gynecol Obstet Invest 44, 265–269 (1997).

47. T. S. Jokiranta, HUS and atypical HUS. Blood 129, 2847–2856 (2017).

48. A. Handler, D. D. Ginty, The mechanosensory neurons of touch and their mechanisms of activation. Nat Rev Neurosci 22, 521–537 (2021).

49. S. S. Ranade et al., Piezo1, a mechanically activated ion channel, is required for vascular development in mice. Proc Natl Acad Sci U S A 111, 10347–10352 (2014).

50. A. T. Chesler et al., The Role of PIEZO2 in Human Mechanosensation. N Engl J Med 375, 1355–1364 (2016).

51. J. Andersen et al., Expression of connexin-43 in human myometrium and leiomyoma. Am J Obstet Gynecol 169, 1266–1276 (1993).

52. A. Conde-Agudelo, R. Romero, J. P. Kusanovic, Nifedipine in the management of preterm labor: a systematic review and metaanalysis. Am J Obstet Gynecol 204, 134 e131–120 (2011).

## References

53. S. H. Woo et al., Piezo2 is required for Merkel-cell mechanotransduction. Nature 509, 622–626 (2014).

54. R. Witschi et al., Hoxb8-Cre mice: A tool for brain-sparing conditional gene deletion. Genesis 48, 596–602 (2010).

55. C. F. Yang et al., Sexually dimorphic neurons in the ventromedial hypothalamus govern mating in both sexes and aggression in males. Cell 153, 896–909 (2013).

56. S. S. Ranade et al., Piezo2 is the major transducer of mechanical forces for touch sensation in mice. Nature 516, 121–125 (2014).

57. S. M. Cahalan et al., Piezo1 links mechanical forces to red blood cell volume. Elife 4, (2015).

58. X. Chen et al., Engineered AAVs for non-invasive gene delivery to rodent and non-human primate nervous systems. Neuron 110, 2242–2257 e2246 (2022).

59. C. C. Rada, S. L. Pierce, C. A. Grotegut, S. K. England, Intrauterine telemetry to measure mouse contractile pressure in vivo. J Vis Exp, e52541 (2015).

60. I. C. Clark et al., Microfluidics-free single-cell genomics with templated emulsification. Nat Biotechnol 41, 1557–1566 (2023).

61. V. Nudell et al., HYBRiD: hydrogel-reinforced DISCO for clearing mammalian bodies. Nat Methods 19, 479–485 (2022).

62. D. R. Stirling et al., CellProfiler 4: improvements in speed, utility and usability. BMC Bioinformatics 22, 433 (2021).

63. S. J. Fleming et al., Unsupervised removal of systematic background noise from droplet-based single-cell experiments using CellBender. Nat Methods 20, 1323–1335 (2023).

64. R. Satija, J. A. Farrell, D. Gennert, A. F. Schier, A. Regev, Spatial reconstruction of single-cell gene expression data. Nat Biotechnol 33, 495–502 (2015).

65. C. S. McGinnis, L. M. Murrow, Z. J. Gartner, DoubletFinder: Doublet Detection in Single-Cell RNA Sequencing Data Using Artificial Nearest Neighbors. Cell Syst 8, 329–337 e324 (2019).

66. I. Korsunsky et al., Fast, sensitive and accurate integration of single-cell data with Harmony. Nat Methods 16, 1289–1296 (2019).

67. B. Phipson et al., propeller: testing for differences in cell type proportions in single cell data. Bioinformatics 38, 4720–4726 (2022).

68. X. Zheng et al., Massively parallel in vivo Perturb-seq reveals cell-type-specific transcriptional networks in cortical development. Cell 187, 3236–3248 e3221 (2024).

